# Is Language a Mechanical Signal? Cytoskeletal Responses to Speech in Yeast

**DOI:** 10.1101/2025.07.08.663738

**Authors:** Mehrta Shirzadian, Emanuel Gollob, Christoph Reiter, Ulla Rauter, Manuel Paschinger, Carolina Caballero, Mark Rinnerthaler, Klaus Spiess

**Affiliations:** Center of Public Health, Medical University Vienna; Department of Biosciences and Medical Biology, University of Salzburg; Creative Robotics, University of Arts Linz; University of Applied Arts, Department Digital Arts, Vienna; University of Music and Performing Arts Vienna

## Abstract

What if vocal language were not only a medium for human communication but a vibrational force that leaves structural traces in living cells? This study explores how audible sound, particularly the structured elements of human speech, affects the cytoskeleton of *Saccharomyces cerevisiae*. Using a direct-contact acoustic system, we exposed yeast to distinct sound types: tonal vibrations, broadband noise, and consonant phonemes. Fluorescence microscopy revealed that tonal stimuli with coherent low-frequency patterns enhanced actin polymerization and shmoo formation, both markers of polarity and mating. In contrast, broadband noise disrupted actin integrity, while consonants produced no measurable effects. These results suggest that rhythmic continuity and spectral coherence, key features of speech, can modulate cytoskeletal organization in non-auditory cells. By reframing vocal language as mechanical input rather than semantic content, this study bridges microbial cell biology with acoustic ecology and proposes a new lens for exploring how human-generated soundscapes physically influence living systems.

## Introduction

Across biological, physical, and cultural domains, sound acts as a pervasive yet understudied form of mechanical energy. This study investigates how audible sound, anging from pure tones to patterned elements of speech, affects the internal structure of microbial eukaryotes. Bridging biophysics, microbiology, and acoustic ecology, we focus on *Saccharomyces cerevisiae* as a model system to explore how cells without auditory organs might still respond structurally to vibrational cues. Our aim is to highlight specific disciplinary gaps: in biophysics, the need for precise acoustic characterization; in microbiology, the overlooked role of the cytoskeleton in sound perception; and in ecoacoustics, the untested influence of linguistic soundscapes on microbial life. Each of the following subsections describes the gap this study addresses within one of these contexts, building toward a cross-disciplinary understanding of how cells might engage with their acoustic environment.

### Sound as a Biophysical Signal

Sound is a mechanical vibration that propagates through air, liquids, or solids, permeating nearly every environment. Traditionally examined through the lens of hearing and psychoacoustics, sound is now increasingly recognized as a physical force capable of influencing biological systems even in the absence of specialized auditory organs [1, 2]. Across organisms from microbes to mammals, cells are equipped with mechanisms to detect and respond to mechanical stimuli such as pressure, shear, and vibration. These stimuli, once transduced into biochemical signals, can modulate processes like growth, adhesion, metabolism, and differentiation [2, 3].

In microbial organisms that lack dedicated sensory structures, acoustic signals can still impact cells indirectly through vibrational energy transferred from their surrounding environment. Oscillatory displacements, traveling through liquids, gels, or container walls, can reach cellular membranes and intracellular structures. For instance, *Escherichia coli* increases biomass yield and stress resilience when exposed to tones around 5 kHz [4], and *Saccharomyces cerevisiae* shows altered metabolism and growth under specific acoustic conditions [5]. To systematically summarize these diverse microbial responses to audible frequencies, we compiled data from multiple studies, categorizing responses into growth stimulation, inhibition, or significant metabolic shifts (Figure 1). This synthesis highlights frequency ranges (100–1000 Hz and 1.5–2.2 kHz) consistently reported to promote microbial growth across various organisms. Frequencies outside these optimal windows tend to yield neutral or inhibitory outcomes, emphasizing the nuanced relationship between acoustic parameters and biological effects.

**Figure 1:**
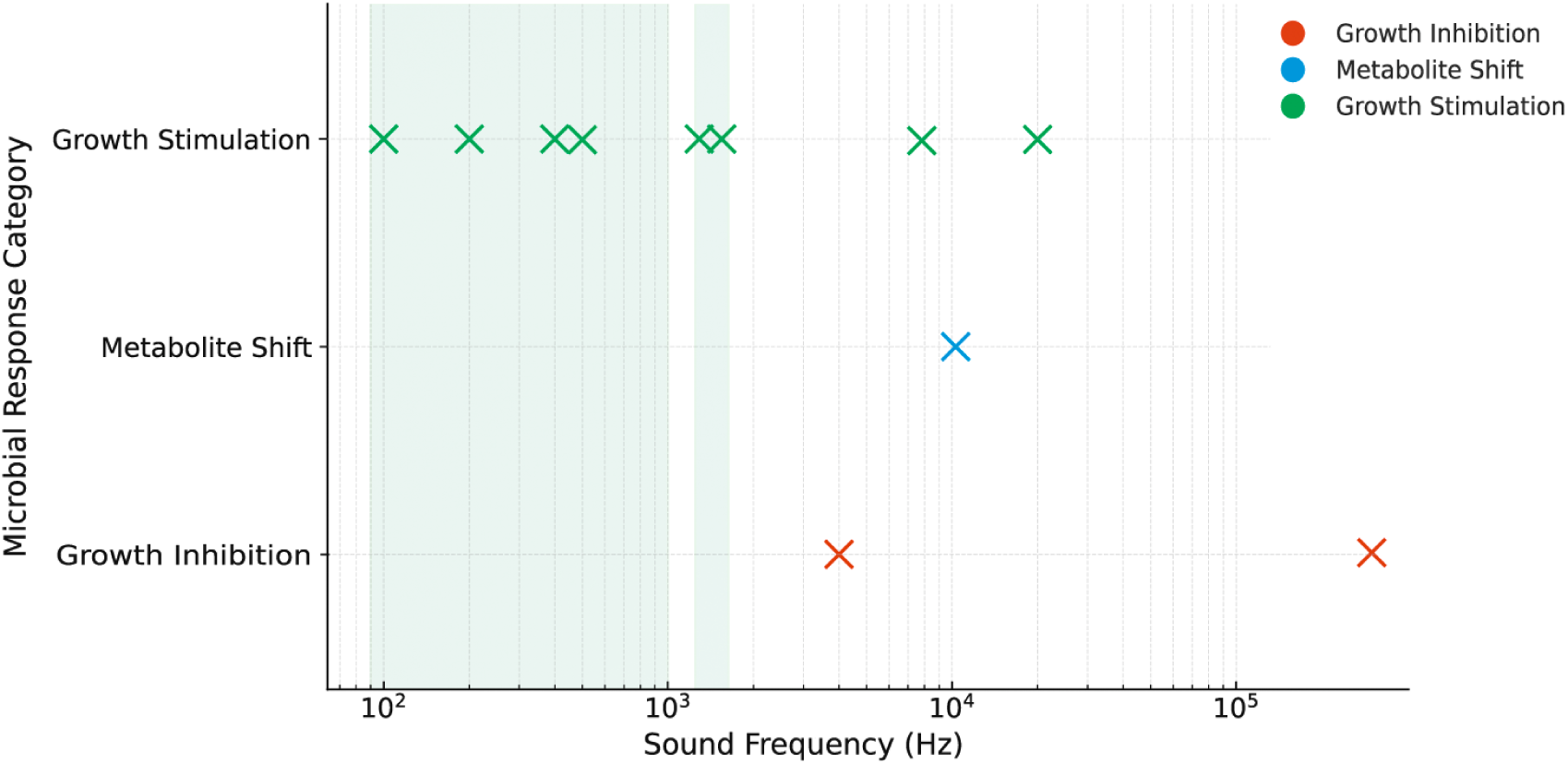
Mapped sound frequencies vs. microbial response categories (compiled from multiple studies). Green points denote growth stimulation (increased growth rate or biomass); red points denote growth inhibition (decreased growth or viability); the blue point indicates a notable metabolic shift (significant biochemical changes with minimal net growth gain). Shaded green bands highlight consensus frequency ranges (100–1000 Hz and 1.5–2.2 kHz) where multiple studies report growth promotion [6–9]. Frequencies above these ranges tend to produce neutral or inhibitory effects in many bacteria, whereas mid-to-low audible frequencies (hundreds of Hz to 1 kHz) repeatedly show positive growth outcomes across diverse single-species microbes.

Building on these findings, several recent studies have explored how sound may influence yeast metabolism and fermentation in context-dependent ways. Harris et al. [10] found that audible frequencies increased yeast growth and altered volatile profiles. Similarly, Adadi et al. [11, 12] reported that sound exposure accelerated fermentation and slightly shifted aroma compound production. However, Benítez et al. [13] showed minimal effects when sound was delivered through liquid, highlighting the importance of stimulus modality. These findings suggest a potential role for acoustic stimulation in optimizing fermentation processes, while also underscoring the importance of how sound is delivered to biological systems.

Despite a growing body of evidence suggesting sound may influence biological systems, the reproducibility of such findings remains inconsistent. A 2023 study by Benitez et al. [14] examined the impact of audible sound on *S. cerevisiae* fermentation using a tightly controlled setup. When experiments were conducted in an anechoic chamber and sound was delivered directly into the liquid culture to avoid environmental artifacts, the team observed little to no biological change. These null results contradicted earlier reports and led the authors to suggest that prior effects might have arisen from uncontrolled factors for example, vibrations traveling through the air and coupling into flask walls, or local heating, convection, and fluid agitation. Notably, when Benitez and colleagues used submerged sound sources to eliminate external vibrational interference, biological changes largely disappeared [14].

Similarly, while some studies report stimulation of growth, migration, or enzymatic activity under sound exposure, others reveal no effect or even inhibition. Such variability likely stems from differences in experimental setups: frequency range, waveform, sound pressure level, exposure duration, and apparatus geometry all play critical roles in shaping biological outcomes [15]. It remains unclear whether the observed effects arise from direct mechanosensation, or from secondary physical processes such as microthermal gradients, altered gas exchange, or mechanical shear. These inconsistencies point to a critical gap: the **lack of standardized, biologically meaningful sound characterization**. Without consistent terminology, reproducible acoustic parameters, and controlled delivery methods, comparison across studies is difficult and mechanistic insights remain elusive.

### From Cymatics to Cytoskeletal Mechanotransduction

Despite the growing number of studies reporting biological effects of audible sound on microbes, the underlying mechanisms remain poorly characterized. While empirical evidence points to measurable changes in growth, metabolism, and morphology across species, there is currently no unified theory to explain how mechanical vibrations elicit such responses in organisms that lack auditory systems. Most existing research is phenomenological rather than mechanistic, focusing on observed outcomes rather than the pathways involved. This conceptual gap has led some to draw on analogies such as cymatics. Classic cymatics demonstrations, such as particles forming intricate shapes on vibrating surfaces, are often cited as metaphors for the organizing potential of sound in biology [16]. However, applying this metaphor to living systems requires caution. Cells are not passive particles but active, self-organizing systems that continuously remodel their internal structures through energy-dependent processes. As such, they are more likely to exhibit transient, regulated responses to mechanical inputs than to undergo direct geometric patterning. Claims of sound imprinting stable structures on living tissues have thus been criticized for lacking mechanistic clarity [17].

Recent studies, however, have begun to reveal how cells may actively perceive and respond to vibrational cues. In mammalian systems, sound exposure has been shown to affect cytoskeletal organization, gene expression, and rhythmic activity. For instance, HL-1 cardiac cells exposed to specific sound frequencies exhibit distinct actin and microtubule architectures, indicating that cells can differentiate among mechanical signals [18]. These results suggest that the cytoskeleton acts not just as a structural framework but as a sensor of physical stimuli, converting vibrational input into biochemical and structural outcomes [19].

Despite growing recognition of these effects in animal systems, the role of the cytoskeleton in microbial responses to sound remains largely unexamined. Studies on acoustic stimulation in microbes, including bacteria and yeast, have typically focused on growth rates, metabolic outputs, or stress resistance, rarely considering the structural consequences of mechanical vibration. This is a significant omission in the case of *Saccharomyces cerevisiae*, a model eukaryote with a well-characterized and highly responsive actin cytoskeleton.

One particularly relevant yet understudied morphological phenomenon is shmoo formation: the polarized outgrowth yeast cells develop in response to mating pheromones. Shmooing requires precise actin remodeling and represents a clear, quantifiable readout of cytoskeletal dynamics [20, 21]. Despite its clarity as a structural marker, shmoo formation has not previously been investigated in the context of acoustic stimulation. Whether sound can modulate or mimic the polarization behavior typically triggered by pheromones remains unknown. This gap highlights a broader need to examine how vibrational inputs affect cytoskeletal morphology in microbial systems and whether observable outcomes like shmooing could serve as reliable indicators of mechanoresponsive behavior.

### Language as Acoustic Ecology: A Bridge Between Culture and Microbial Life

Emerging research in ecoacoustics and sonobiology suggests that acoustic environments can shape microbial life: microorganisms have shown sensitivity to various frequencies, adjusting growth, community composition, and metabolic outputs in response to sonic stimuli [1, 22, 23]. While most studies have employed generic tones, the specific effects of linguistic sound-scapes, the patterned, rhythmic structure of spoken language, have not been systematically investigated.

To situate this inquiry, we briefly clarify two key linguistic concepts. *Semantics* is the study of meaning in language, how words and utterances represent concepts, intentions, and referents in the world [24]. *Phonetics*, by contrast, examines the physical properties of speech sounds: how they are produced, transmitted as vibrations in the air, and perceived by listeners [25]. In natural language, these domains typically remain distinct; however, certain phenomena, such as *onomatopoeia*, words that phonetically imitate the sounds they describe (e.g., *buzz*, *clang*, *hiss*), provide a direct bridge between phonetics and semantics [26]. In onomatopoeia, the vibrational signature of a word conveys semantic meaning not arbitrarily, but through acoustic resemblance.

Critically, linguistic soundscapes are composed of systematic acoustic patterns. Speech sounds have measurable *frequencies* (cycles per second, Hz) that contribute to pitch [27]. *Periodicity* refers to the regular repetition of waveforms, especially in voiced speech sounds [27], while *rhythm* describes the temporal structuring of syllables, stress, and intonation patterns across time [28]. These patterned acoustic features, not just raw sound energy, make up the vibratory structure of language.

This study proposes that such structured speech, may influence microbial behavior in ways more nuanced than previously considered. By framing language as a form of patterned acoustic energy, this study invites a new perspective: human culture as a mechanical participant in microbial ecologies. This conceptual bridge between structural biology and cultural sound reframes language not only as a medium of meaning, but also as an environmental force. It opens the question: what biological effects might emerge from the patterned vibrational presence of human beings in the world?

### Objective and Broader Implications

This study investigates whether audible sound, delivered in a physically controlled and biologically meaningful way, affects the actin cytoskeleton and shmoo formation in *Saccharomyces cerevisiae*. Using this well-characterized model and defined acoustic parameters, we aimed to generate reproducible results that could serve as a foundation for future comparisons. Rather than focusing on broad physiological outcomes, we examined the cytoskeleton and associated shmoo morphology as mechanosensitive structures that reflect how cells perceive and respond to physical cues. We hypothesize that specific sound frequencies, including components of human speech, may influence cytoskeletal organization and cellular polarity, suggesting that language might function as a mechanical input at the cellular level.

To explore this, we first tested tonal sounds and broadband noise as acoustically distinct proxies for speech components: tones approximate the harmonic structure of vowels, while noise resembles the irregular, broadband character of many consonants. These contrasts provided a starting point for evaluating the cytoskeletal and shmooing responses to vibrational input. We then tested isolated consonants to examine more complex, discontinuous acoustic structures. While these transient sounds did not appear to elicit detectable cytoskeletal or morphological changes, their inclusion helps refine how future studies might explore language as a vibrational force. Fragmented, brief stimuli such as individual consonants may be too short or acoustically variable to produce measurable structural effects. This underscores the importance of continuity, rhythm, and acoustic density when designing experiments on speech–microbe interactions. Taken together, this work outlines a methodological foundation for studying how the structured mechanical energy of language might modulate cellular architecture and polarization.

## Methods

## Results

### Tonal Sound Enhances Cytoskeletal Polarization and Shmooing

Cells exposed to the tonal stimulus showed visibly enhanced actin polymerization and cytoskeletal organization. Fluorescence imaging revealed stronger actin intensity and increased cortical coverage (Figure 2A–C). Most notably, cells under tonal conditions frequently exhibited shmoo formation, a mating-associated morphological change that requires actin remodeling. This morphological change is also evident in representative brightfield images (Figure B), where cells exposed to tonal vibration display elongated projections characteristic of mating readiness. These effects suggest that coherent low-frequency vibrations can promote polarity and mating-like states in yeast.

**Figure 2:**
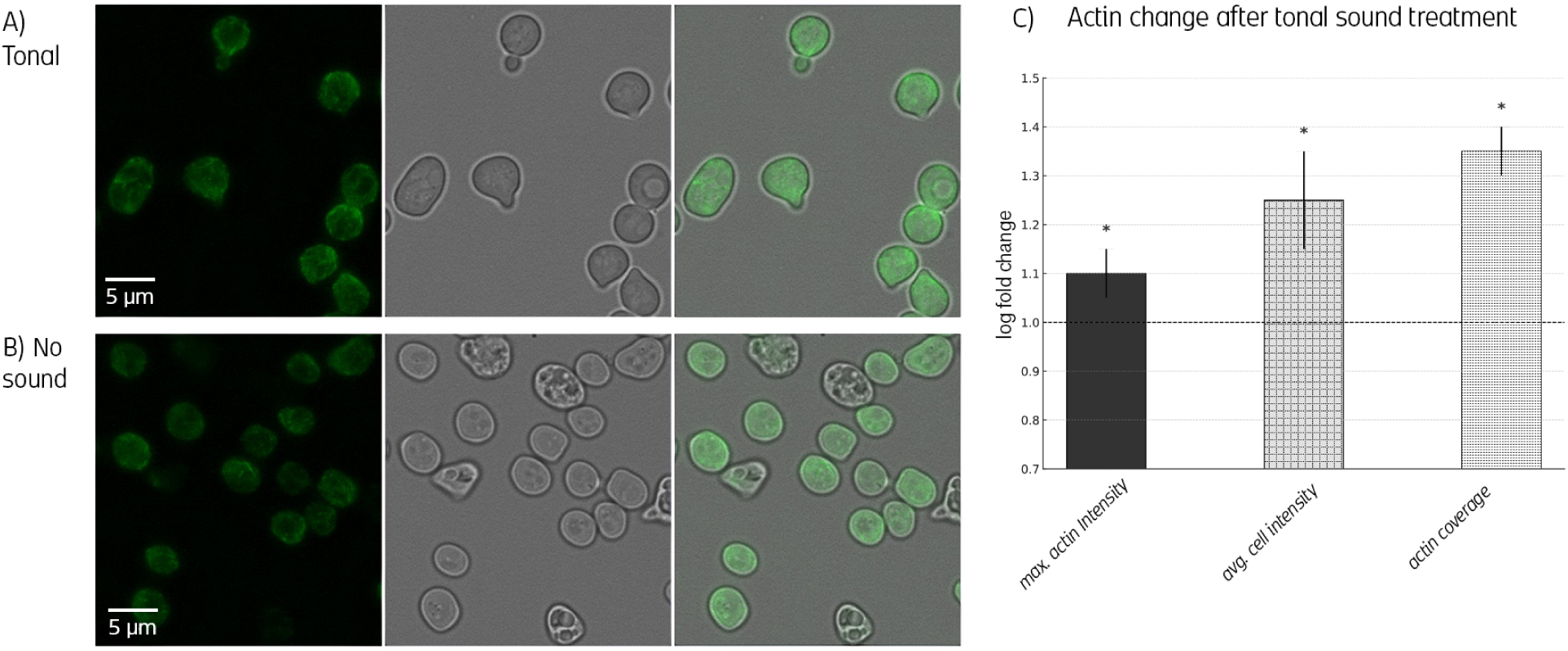
Tonal vibration enhances cytoskeletal polarity, actin intensity, and morphological differentiation. **(A)** Fluorescence microscopy images show increased actin signal in cells exposed to tonal sound, including clear cortical localization and frequent formation of shmoo projections, hallmarks of polarity and mating readiness. **(B)** Segmentation overlays reveal actin accumulation along polarized sites and morphological asymmetry consistent with cytoskeletal remodeling. **(C)** Bar plots quantify the effect of tonal stimulation on cytoskeletal metrics. Maximal actin intensity, average cellular intensity, and actin coverage are significantly elevated relative to no-sound controls (log fold change > 1, p < 0.05). This indicates that coherent, low-frequency vibration can amplify structural readiness for morphogenesis and promote polarity-driven behaviors.

### Noise Disrupts Cytoskeletal Organization and Inhibits Shmooing

In contrast, broadband noise exposure led to a marked reduction in actin polymerization. Cells exhibited diminished actin signal, reduced cortical organization, and less defined polarity. These inhibitory effects were consistently observed across replicates and point to a form of vibrational stress (Figure 3).

**Figure 3:**
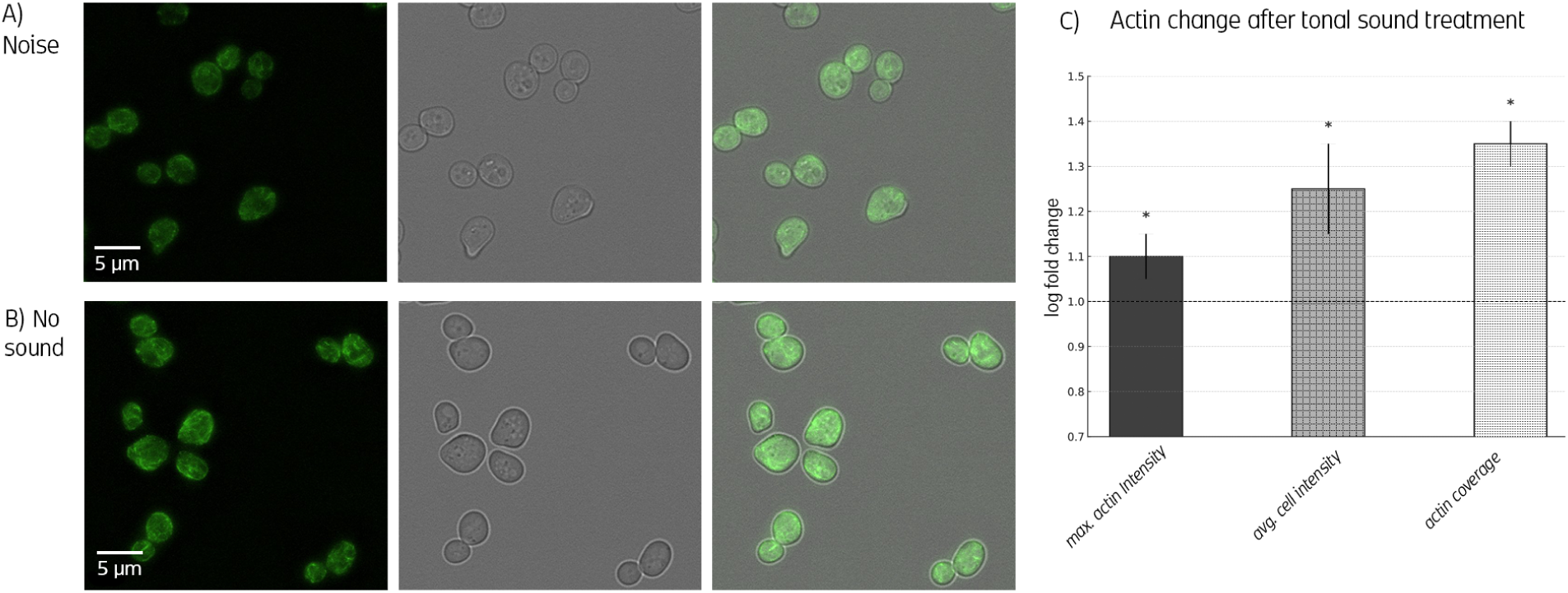
Aperiodic broadband noise suppresses actin organization, intensity, and polarity cues. **(A)** Fluorescence microscopy of noise-treated cells shows diffuse, low-intensity actin signal with no visible polarization or shmoo formation. **(B)** Segmentation confirms the absence of cortical actin enrichment or asymmetric morphologies, reflecting cytoskeletal disorganization. **(C)** Quantification reveals a significant reduction in maximal actin intensity and average actin levels (p < 0.05 and p < 0.001, respectively), while total actin coverage remains unchanged or trends downward. These results suggest that disordered acoustic input acts as a mechanical stressor, inhibiting polarity and destabilizing cytoskeletal coherence.

In parallel, the disordered, high-frequency acoustic energy appeared to interfere with shmoo formation, preventing the emergence of polarized, mating-like morphology. This suppression is clearly visible in Figure B, where cells exposed to noise lack the asymmetric projections characteristic of tonal stimulation.

### Consonant Stimuli Show Minimal or Inconsistent Effects

Despite acoustic similarities to noise, the consonant stimulus had no observable impact on actin organization or shmooing. Neither intensity nor morphological markers differed from control. This may reflect the acoustic heterogeneity of consonants—especially their temporal discontinuity and abruptness which could prevent sustained mechanical engagement with cells. The brief, fragmented structure of the consonant signal likely limits its capacity to trigger structural responses in yeast.

### Summary of Cellular Responses Across Sound Conditions

A consolidated comparison across conditions revealed a clear dichotomy: tonal sound enhanced cytoskeletal dynamics and morphological differentiation, while noise disrupted both. Consonants produced no consistent effect. These findings suggest that not only frequency content but also temporal coherence and rhythmic continuity may be critical in modulating cellular responses to sound. (Figure 4)

**Figure 4:**
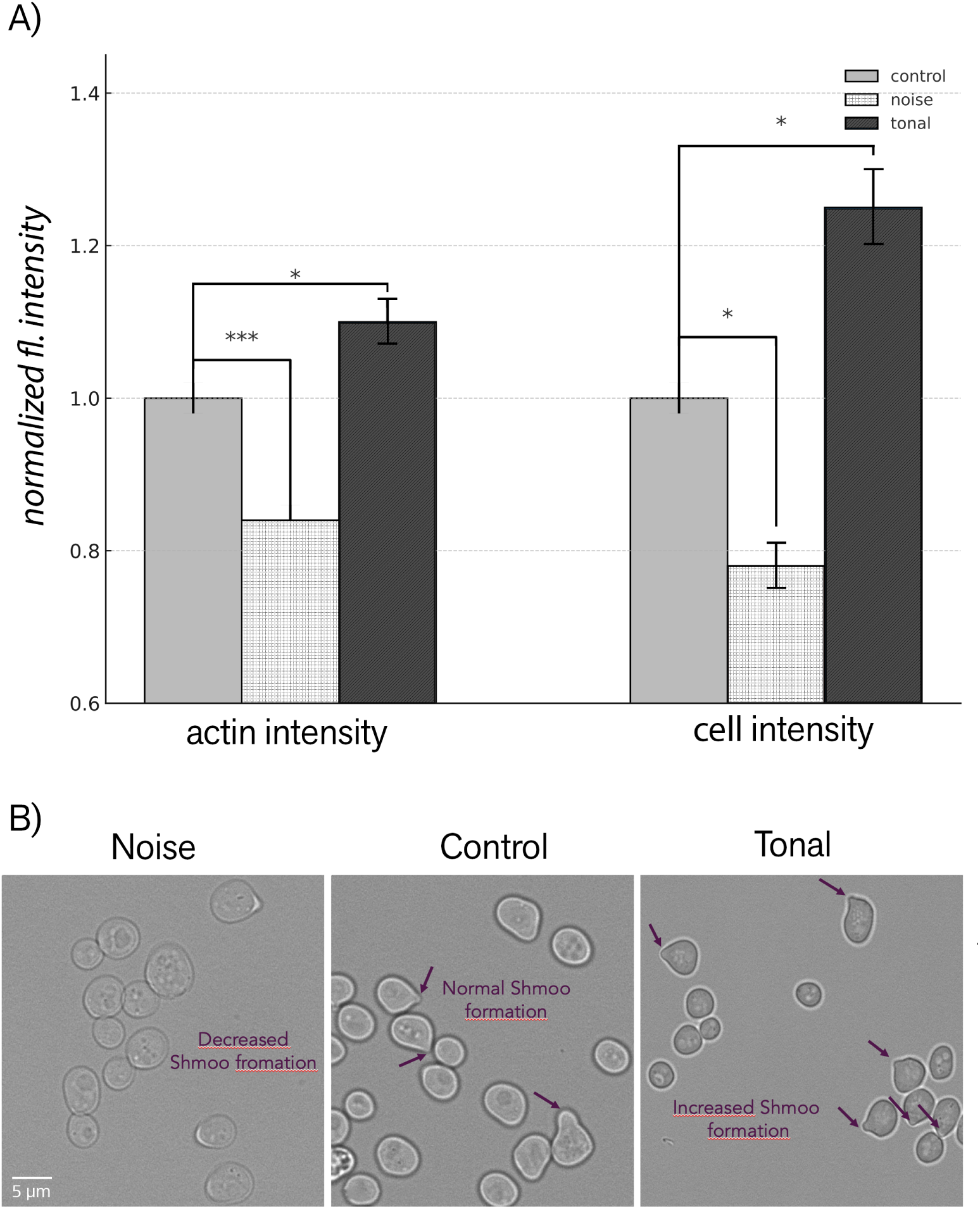
Tonal sound enhances, and broadband noise suppresses, actin and cell fluorescence intensities. **(A)**Bar plot summarizing normalized fluorescence intensity measurements across conditions. Actin intensity and overall cellular fluorescence intensity were elevated under tonal stimulation and suppressed under broadband noise, compared to control. Values are normalized to the no-sound control within each replicate (set to 1) and reflect fold changes. Statistical significance was assessed via one-sample t-tests with Ben-jamini–Hochberg FDR correction (n *≥* 3). Error bars indicate standard deviation. Asterisks denote statistical significance: p < 0.05. **(B)** Comparative brightfield images of yeast morphology under different sound conditions. Cells exposed to tonal sound exhibit pronounced shmoo formation, indicative of cytoskeletal polarization. Control cells show standard budding morphology, while cells under broadband noise display reduced or absent shmooing.

## Discussion

This study demonstrates that audible sound can elicit structural and morphological changes in yeast cells, depending on the acoustic characteristics of the stimulus. Using fluorescence microscopy to visualize actin filaments, we observed that tonal sound promoted actin polymerization and shmoo formation, while broadband noise suppressed both features. Consonant-rich stimuli, in contrast, showed no measurable impact on cytoskeletal organization.These distinct outcomes suggest that frequency content alone is not sufficient to predict cellular response; rather, the temporal coherence and spectral continuity of the stimulus appear to play a central role.

Most previous studies have focused primarily on the role of frequency in shaping microbial responses to sound. In contrast, our findings emphasize that temporal structure and spectral continuity, features often overlooked, may be critical determinants of biological relevance.

The ability of pure tonal sound to promote cytoskeletal polarization aligns with prior work on mechanical signaling in eukaryotic cells [18]. In our case, enhanced actin intensity and increased shmoo incidence under tonal stimulation suggest that rhythmic, low-frequency vibrations may mimic aspects of mating pheromone signaling, possibly by reinforcing cytoskeletal alignment or endocytic trafficking pathways necessary for projection formation.

By contrast, broadband noise appeared to disrupt cytoskeletal integrity, reducing actin signal intensity and suppressing morphological differentiation. This mirrors previous observations of vibrational noise acting as a stressor in microbial systems [15], possibly through random shear forces or inconsistent mechanical inputs that interfere with cellular organization. Such noise-induced suppression may result from either cytoskeletal disassembly or global energy reallocation under stress-like conditions.

Although this study examined cytoskeletal morphology rather than cell viability markers, it is noteworthy that perturbations in actin organization, such as the depolarization of actin cables or disruption of cortical patches, have been associated with apoptotic signaling in yeast and are thought to reflect conserved stress responses across eukaryotes [29–31]. Stabilization or dysregulation of filamentous actin (F-actin) through genetic manipulation or actin-stabilizing agents has been shown to induce mitochondrial dysfunction, reactive oxygen species (ROS) accumulation, and cell death in *Saccharomyces cerevisiae* [30, 31]. In this context, the disassembly of the actin cytoskeleton observed under broadband noise exposure could reflect the initiation of stress responses, potentially including programmed cell death pathways. Supporting this interpretation, recent work has demonstrated that defects in actin dynamics activate stress MAPK pathways that promote the elimination of cytoskeletally compromised yeast cells [32]. However, since apoptosis was not directly assayed in our experiments, this remains a speculative interpretation. Future studies employing canonical apoptotic markers, such as ROS accumulation, phosphatidylserine externalization, and caspase-like activity (metacaspase) will be necessary to determine whether sound-induced actin disruption correlates with programmed cell death [31].

Interestingly, consonant stimuli, though partially overlapping with noise in their acoustic profile, produced no discernible biological effects. This may stem from their temporal fragmentation: consonants are characterized by brief, transient bursts of energy that lack the sustained waveform required to activate mechanotransductive pathways. While some consonants (e.g., fricatives) showed high zero-crossing rates and spectral centroids, the lack of continuity may prevent cells from integrating these signals over time. These findings indicate that duration and rhythmic structure, not just spectral features, critically determine whether a sound functions as a biologically relevant mechanical input.

Beyond cytoskeletal effects, our results have important implications for interpreting optical density (OD_600_) measurements during audio stimulation experiments. Shmoo formation involves morphological elongation and growth arrest, both of which alter the optical scattering properties of the cells. As cells become polarized and begin to aggregate due to mating-associated adhesion proteins, they scatter light less efficiently per unit biomass [**lipke2023fungal**, 33]. This can result in reduced OD readings even when total cell number or biomass remains constant. Additionally, clumped or sedimented cells may escape detection by OD altogether. Therefore, a decline in OD during tonal sound exposure, as observed in pilot studies, should not be interpreted as cell death or growth inhibition, but rather as a manifestation of morphological transition and aggregation. This reinforces the need to supplement OD measurements with microscopy or direct cell counts in vibration-sensitive conditions.

Taken together, our findings suggest that mechanical signals encoded in sound, especially rhythmic, structured vibrations, can modulate cytoskeletal dynamics and polarity in microbial cells. While the precise biophysical mechanisms remain to be elucidated, possible pathways include membrane tension sensing, actin–membrane coupling, or modulation of trafficking and endocytosis. Our observation that consonants had no effect further implies that cells may require prolonged vibrational coherence to mount a structural response.

Future directions include embedding consonants in full linguistic phrases, testing environmental language exposure in multispecies communities, and investigating downstream transcriptional changes following mechanoacoustic stimulation. This study positions language not only as a cultural artifact but also as a mechanical force with potential biological consequences, opening a novel perspective on how communication leaves molecular traces in shared environments.

## Limitations of the Study

This study used a specific yeast strain and experimental setup to investigate cytoskeletal responses to sound stimuli. While our direct-contact stimulation approach improves mechanical signal precision, it does not capture the full diversity of acoustic environments cells may encounter in nature. Moreover, we used only one fluorescent marker (ABP140-GFP), so broader cytoskeletal or transcriptomic changes remain to be explored. Future studies could test other strains, mating types, or vibration parameters to confirm and generalize our findings.

## Acknowledgments

This work was supported by the Austrian Science Fund (FWF), project AR 687. We thank all contributors and collaborators who supported this study. We also thank Hanna Hofmann for her graphic design support.

## Author Contributions

M.S. conducted the literature review, created visualizations, and wrote the main manuscript text. E.G. designed and implemented the acoustic stimulation system and contributed to content development. C.R. performed data analysis, generated plots, and contributed to writing the Methods section. U.R. generated and designed the sound files and contributed to the experimental procedures. M.P. carried out laboratory experiments and performed microscopy image acquisition. C.C. conducted acoustic analysis of the sound stimuli and contributed to the preparation of one figure. M.R. supervised the microbiological work, image acquisition, and data analysis. K.S. developed the conceptual framework of the study, supervised the project, and edited the manuscript. K.S., M.S., C.R., and M.R. reviewed and approved the final version of the manuscript.

## STAR⋆Methods

### Key Resources Table

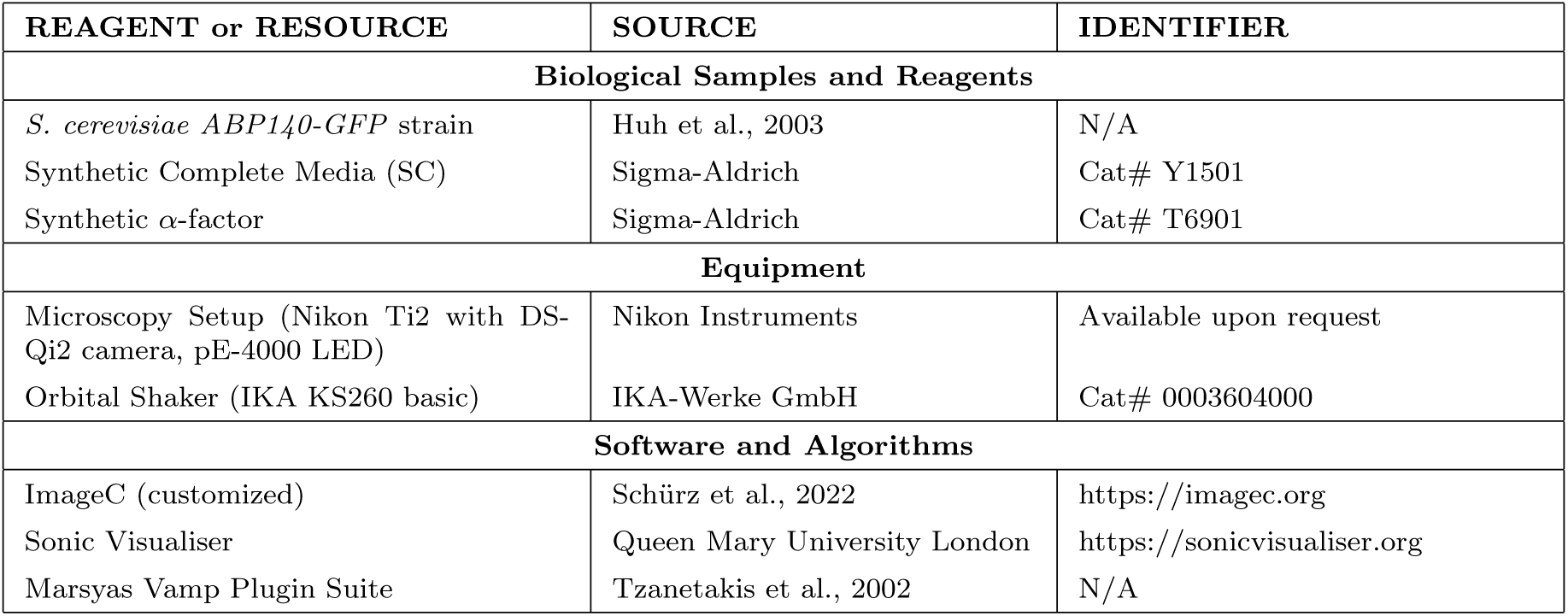

| REAGENT or RESOURCE | SOURCE | IDENTIFIER |
| --- | --- | --- |
| **Biological Samples and Reagents** |  |  |
| *S. cerevisiae* ABP140-GFP strain | Huh et al., 2003 | N/A |
| Synthetic Complete Media (SC) | Sigma-Aldrich | Cat# Y1501 |
| Synthetic α-factor | Sigma-Aldrich | Cat# T6901 |
| **Equipment** |  |  |
| Microscopy Setup (Nikon Ti2 with DS-Qi2 camera, pE-4000 LED) | Nikon Instruments | Available upon request |
| Orbital Shaker (IKA KS260 basic) | IKA-Werke GmbH | Cat# 0003604000 |
| **Software and Algorithms** |  |  |
| ImageC (customized) | Schürz et al., 2022 | https://imagec.org |
| Sonic Visualiser | Queen Mary University London | https://sonicvisualiser.org |
| Marsyas Vamp Plugin Suite | Tzanetakis et al., 2002 | N/A |

**Note:** Additional materials and data are available upon request from the Lead Contact.

### Lead Contact

Further information and requests for resources, materials, or data should be directed to the Lead Contact, Klaus Spiess.

### Materials Availability

This study did not generate new unique biological materials. Microscopy setups and sound exposure devices are available upon reasonable request from the Lead Contact.

### Sound Exposure Setup

Acoustic stimulation was delivered using a custom-built apparatus in which piezoelectric transducers were directly affixed to the outer surface of culture flasks, allowing transmission of mechanical vibrations into the liquid without the need for airborne propagation. Audio files were digitally generated and amplified using a compact audio amplifier connected to the transducers.

Five experimental sound conditions were tested:

- NS = No sound (control)
- 100 = Pure tone at 250 Hz
- 0 = High-pass filtered noise (>2000 Hz)
- V = Human speech vowels
- C = Human speech consonants

Stimulation intensity was standardized to approximately 80 dB SPL at the transducer-liquid interface, verified using a contact vibration sensor. All exposures were performed under static incubation at 28*^◦^*C for 1.5–2 hours.

This direct-contact approach represents a significant improvement over conventional speaker-based systems, which often suffer from variability due to room acoustics and spatial inhomogeneity. By contrast, our system ensures reproducible, localized mechanical perturbations. Although prior studies have exposed yeast to airborne music or tones [34], this methodology provides higher mechanical fidelity and minimal signal loss.

### Yeast Strain and Growth Conditions

For the analysis of the actin cytoskeleton in dependence on sound exposure, a yeast strain from the Hu Yeast GFP collection was used [36]. This strain collection, based on yeast strain BY4741 with genotype *MATa his3*Δ*1 leu2*Δ*0 met15*Δ*0 ura3*Δ*0*, includes 4159 yeast strains, each expressing a different full-length ORF fused to a GFP tag.

The strain used in this study expresses *ABP140-GFP*, an AdoMet-dependent tRNA methyltransferase. This protein has an actin-binding domain at the N-terminus and labels actin cables (non-branched actin) and patches (branched actin) in yeast [37]. Actin itself cannot be tagged with GFP, as this is lethal under our experimental conditions.

Cells were grown in synthetic media composed of 2% (w/v) D-glucose, 0.17% (w/v) yeast nitrogen base without amino acids, 0.5% ammonium sulfate, and 10 mL of complete dropout mixture per liter (0.2% Arg, 0.1% His, 0.6% Ile, 0.6% Leu, 0.4% Lys, 0.1% Met, 0.6% Phe, 0.5% Thr, 0.4% Trp, 0.1% Ade, 0.4% Ura, 0.5% Tyr).

An overnight culture was diluted to an OD_600_ of 0.1 and grown under constant shaking until the exponential growth phase was reached. To explore interactions between acoustic stimulation and mating pathway signaling, the *ABP140-GFP*-tagged *MATa* strain was treated with synthetic α-factor, a yeast pheromone. Treatments were applied during early exponential phase using concentrations of 0, 2.8, 5, or 10 µM, with 5 µM used in most experiments. Pheromone was administered alone or simultaneously with sound exposure. Cells were then incubated under static conditions at 28 °C for 1.5–2 hours, a timeframe sufficient to induce shmoo formation and actin polarization.

### Microscopy and Image Analysis

Microscopy was performed using a Nikon Ti2 widefield microscope equipped with a Plan Apo λ 100*×* oil immersion objective (NA 1.45) and a Nikon DS-Qi2 monochrome camera (14-bit), controlled via NIS-Elements AR software (v5.40.00).

For fluorescence imaging, excitation and emission were provided by a CoolLED pE-4000 light source and a FITC filter set (Ex 470 nm / Em 514 nm; filter: 558 nm). Z-stacks of 20 planes were acquired at 0.2 µm step size, with a voxel size of 0.0735 *×* 0.0735 *×* 0.2 µm^3^. Images were captured with 300 ms exposure time, gain 57.7*×*, 1*×*1 binning, and internal triggering.

For brightfield imaging, illumination was provided via the Ti2 transmitted light path (DIA) with an iris intensity of 67.4%. Images were acquired with a 1 ms exposure time, gain 64*×*, and 1*×*1 binning, at a pixel size of 0.0735 *×* 0.0735 µm^2^. Image files followed a structured naming convention:

(runtime in minutes)-(sound condition)-(pheromone concentration in µM)-(FL or DL)-(image index).nd2

Example: 150-NS-2.8-DL-2.nd2 indicates 150 minutes, no sound, 2.8 µM-factor, differential light (DL), image 2.

Fluorescence microscopy images were analyzed using *ImageC*, a powerful AI-assisted software originally developed for the detection of fluorescently labeled extracellular vesicles [35]. Due to its modular and adaptable architecture, the tool was successfully customized for analysis of both brightfield and fluorescence microscopy images of *S. cerevisiae*. Using the brightfield channel, an automated cell recognition pipeline was established, extracting parameters such as cell count and cell area. These segmented cells served as precise masks for evaluating cytoskeletal organization in the fluorescence channel.

The fluorescence analysis pipeline included background correction (rolling-ball subtraction), edge detection, and binary thresholding to segment actin-rich regions. Quantitative metrics extracted per cell included average actin intensity, maximum actin intensity (highlighting polarity hotspots), and total actin coverage (area fraction per cell). Only actin signals within the identified cell boundaries were measured, ensuring robust exclusion of background fluorescence and maintaining data consistency—even in datasets with low signal-to-noise ratios due to genomic GFP expression.

While this study employed only a single fluorescence channel, ImageC supports multichannel datasets and complex fluorophore cross-correlation. Once optimized using a small test set, the workflow allowed high-throughput processing of experimental images. Final results were manually verified and exported as Excel tables for downstream statistical analysis. The user-friendly interface, along with dedicated developer support, enabled efficient deployment of this adaptable, feedback-driven platform. Detection thresholds and segmentation settings were fine-tuned as needed to accommodate differences in illumination and fluorophore intensity across image sets.

### Shmoo Formation Assessment

Due to limitations in automated detection, shmoo formation was manually quantified. Between 50 and 200 cells per image were scored for presence or absence of mating projections based on established morphological criteria and actin enrichment at projection sites. Shmoo formation percentages were then calculated per treatment.

### Data Normalization and Statistical Analysis

Data from each experiment were normalized relative to the internal no-sound control, set to a baseline value of 1. Statistical significance was determined by one-sample *t*-tests (testing against a hypothetical mean of 1) with Benjamini–Hochberg FDR correction for multiple testing when applicable (n *≥* 3). The significance threshold was set at α = 0.05.

Cell detection relied on a segmentation pipeline designed to identify yeast morphology and detect shmoo formation based on deviations from normal circularity. Since shmoos introduce prominent shape asymmetry, this method offered a reliable proxy for quantifying pheromone-induced polarization across treatment conditions.

Beyond morphological classification, quantitative metrics included the percentage of cells displaying shmoos, actin signal intensity, and percent area coverage of the actin signal. Low signal-to-noise ratios in some fluorescence datasets required fine-tuning of intensity thresholds, particularly where signal strength varied between replicates. Consistency in microscopy acquisition settings across experiments was critical to enable cross-sample comparison.

### Acoustic Feature Analysis

To characterize the acoustic properties of the stimuli used in the experiments, audio files were analyzed using *Sonic Visualiser* in combination with the Marsyas plugin suite [38, 39]. This open-source toolchain was selected for its precision in extracting low-level signal descriptors and its reproducibility in experimental acoustic analysis.

Three key acoustic features were extracted from each sound condition (tonal, consonants, vowels, and noise):

1. **Spectral Centroid:** Represents the “center of mass” of the frequency spectrum and correlates with perceived brightness or pitch. It was calculated from the frequency-weighted average of the power spectrum.
2. **Spectral Flatness:** Measures how noise-like versus tone-like a signal is, defined as the ratio of the geometric mean to the arithmetic mean of the power spectrum. Values close to 0 indicate peaky, tonal spectra; values close to 1 indicate flat, noisy spectra.
3. **Zero-Crossing Rate (ZCR):** Quantifies how often the signal waveform crosses zero amplitude, reflecting temporal irregularity. Low values are characteristic of smooth, periodic signals (tones), while high values indicate turbulence or randomness (noise).

Each audio file was analyzed over its full duration, and mean values for each feature were computed. For consonants, additional segmentation was performed to isolate repeated phonemes (p, k, t) and assess their individual acoustic characteristics. Start and end times for each phoneme segment were manually annotated, and the same three features were computed for each.

To better understand how distinct sound inputs might affect cellular responses, we analyzed their spectral and temporal characteristics. The tonal stimulus, while more complex than a pure sine wave, exhibited a highly periodic structure with low spectral flatness and low zero-crossing rate, indicating a smooth, harmonic signal dominated by low-frequency energy. In contrast, the broadband noise signal showed high spectral flatness, high zero-crossing rate, and a bright spectral centroid near 11 kHz, reflecting its turbulent, high-frequency nature.

The consonant stimulus, which included plosives, fricatives, and affricates, showed acoustic features closer to noise, particularly in fricative-rich segments. However, consonants were highly heterogeneous: fricatives like [th] showed much higher spectral centroid and zero-crossing rate than plosives like [p] or [k], suggesting that the overall consonant condition may have been too acoustically fragmented to exert a coherent biophysical influence on cells (Figure 6).

**Figure 5:**
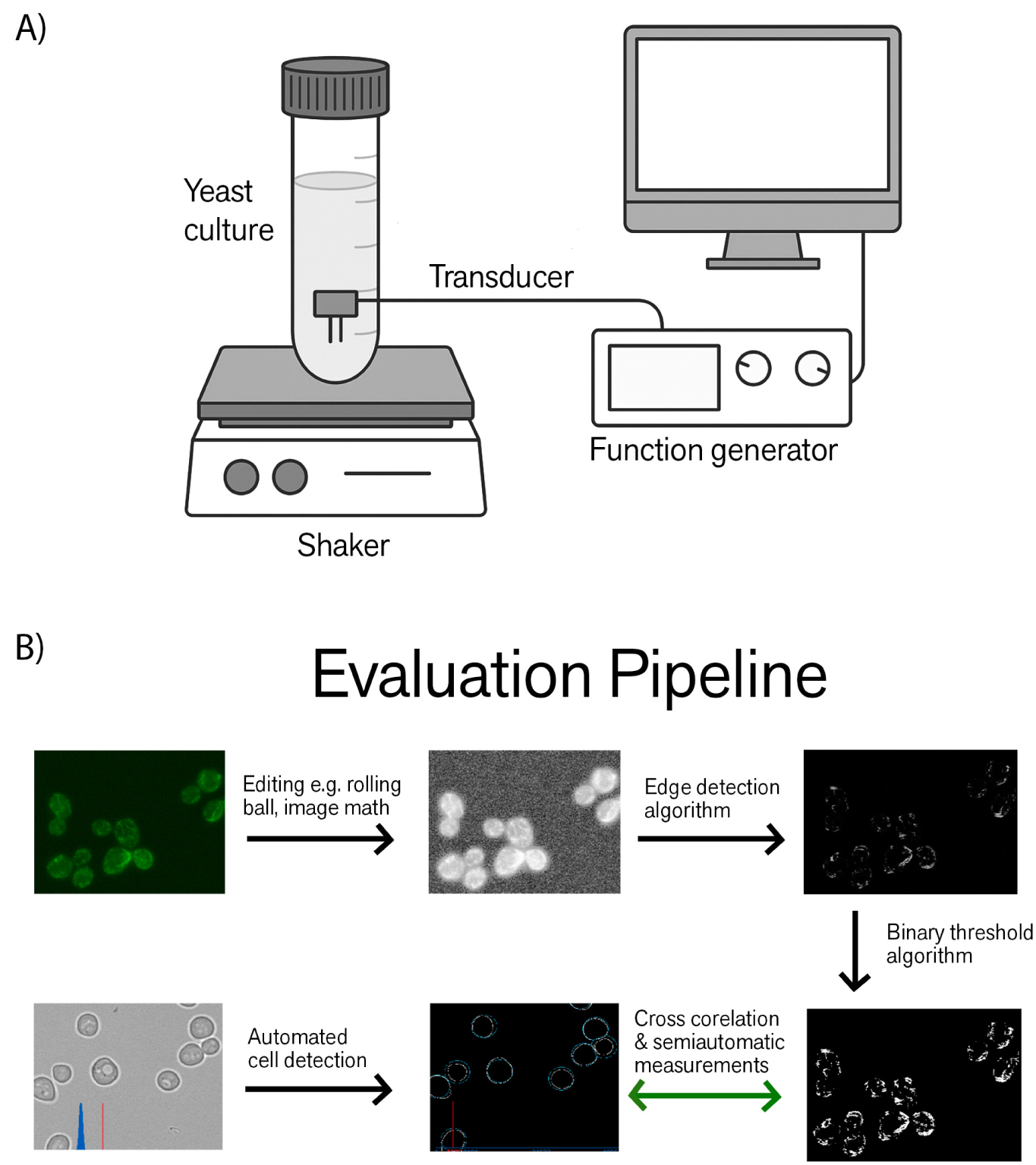
Experimental and evaluation workflow. (A) **Experimental setup for acoustic stimulation.** A small-volume yeast culture is incubated in a sealed test tube mounted on an orbital shaker. A piezoelectric transducer is inserted into the tube via a rubber steglik and connected to a function generator controlled via computer. This setup allows precise delivery of vibrational stimuli directly into the liquid culture, minimizing airborne sound artifacts and enabling reproducible exposure to defined acoustic waveforms. (B) **Evaluation pipeline for cytoskeletal analysis.** Fluorescent and brightfield microscopy images undergo pre-processing (e.g., rolling ball correction), followed by edge detection and binary thresholding to segment cytoskeletal structures. Simultaneously, brightfield images are analyzed for automated cell detection. Cross-correlation and semi-automatic analysis are used to match morphological features to individual cells, allowing measurement of cytoskeletal changes across experimental conditions. Image evaluation was performed using the ImageC software, with support from the developers in adapting the pipeline to our needs [35].

**Figure 6:**
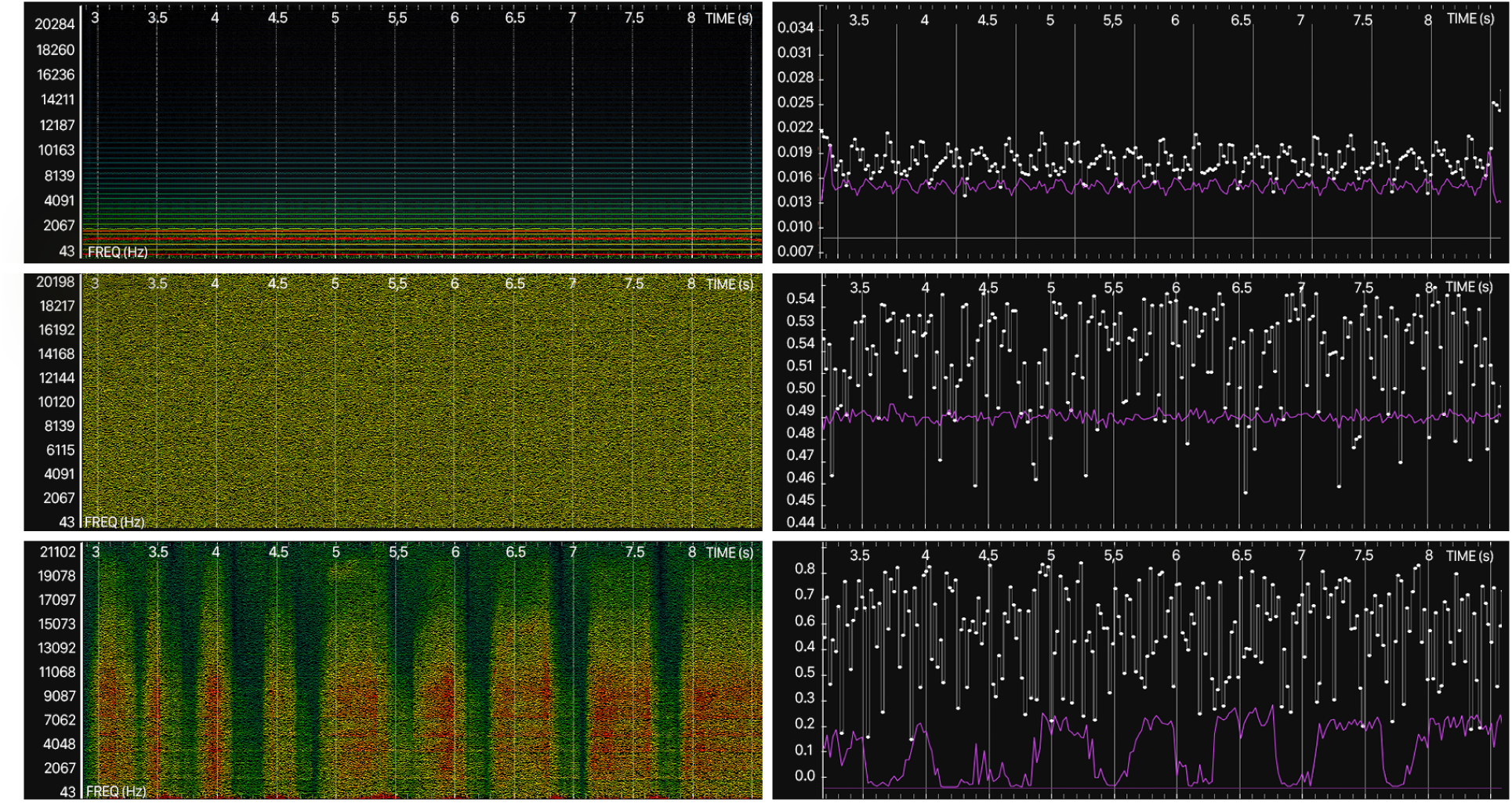
Spectrogram and Audio Feature comparison of acoustic stimuli used in yeast exposure experiments. The figure shows frequency-time representations (left) and plotting (right) of Spectral Centroid (Purple) and Spectral Flatness Values (white) for 5.5 second segments for the three stimuli. (Top) Tonal Sound (Combination of 440, 530 and 1845 Hz), (Middle) Broadband noise (computer generated, uniformly distributed white noise), and (Bottom) Consonant-rich (isolated phonemes). The tonal stimuli shows a stable spectral profile dominated by lower frequencies (under 1000Hz) and harmonic overtones, along with with low Spectral Flatness, consistent with high periodicity. The Broadband noise displays a continuously disordered spectrum with a high frequency Centroid and Spectral Flatness, confirming it’s lack of harmonic structure and it’s highly disordered/noisy energy distribution. The consonant stimulus shows irregular spectral bursts with varied frequency content and relatively high Spectral Flatness, corresponding to the fragmented, noise-like nature of fricative consonants (in the figure, different onsets of the fricative th can be seen).

## Glossary of Terms

Actin cytoskeleton: A dynamic network of actin filaments involved in maintaining cell shape, enabling motility, and organizing intracellular structures. In yeast, it plays a key role in budding and mating projections.
Audible sound: Mechanical vibrations within the frequency range detectable by the human ear (20 Hz to 20 kHz). In this study, it includes pure tones, speech components, and noise.
Broadband noise (noise): An aperiodic acoustic signal comprising a wide and continuous spectrum of frequencies, with no distinct pitch. Characterized by high spectral flatness and zero-crossing rate, it represents mechanical randomness and often functions as a biological stressor.
Consonant: A class of speech sound produced by obstructing airflow, typically brief and rich in high-frequency content. Includes plosives, fricatives, and affricates. Acoustically more fragmented and transient than vowels.
Cymatics: The visual study of how sound vibrations organize matter, often demonstrated through the formation of patterns on a vibrating surface. Used metaphorically to explore how structured vibration may influence biological systems.
Fluorescence microscopy: An imaging technique used to visualize specific structures in cells by tagging them with fluorescent markers. In this study, used to detect actin filament organization and morphological changes.
Mechanotransduction: The conversion of mechanical stimuli into biochemical signals. Key to understanding how cells respond to vibrations or pressure from sound waves.
Phonetics: The scientific study of speech sounds, including their acoustic and articulatory properties. It provides a framework for analyzing the sound structure of language.
Spectral centroid: A measure of the “center of mass” of the frequency spectrum of a signal. Higher values indicate brighter or higher-pitched sounds.
Spectral flatness: A ratio reflecting how noise-like a signal is. Values near 1 indicate uniform energy across frequencies (noisy), while values near 0 suggest tonal, peaked spectra.
Shmoo: A polarized, projection-like morphology formed by yeast during mating. Dependent on actin cytoskeleton rearrangement and used as an indicator of cytoskeletal polarization.
Speech stimulus: A complex acoustic signal composed of structured sequences of phonemes (consonants and vowels), often patterned in time. Used to simulate linguistic acoustic input in experiments.
Tonal sound (tone): A periodic acoustic signal characterized by a narrow frequency range and harmonic structure. Exhibits low spectral flatness and low zero-crossing rate. In this study, used to simulate vowel-like structure.
Vibration: A physical oscillation or repetitive motion, such as those produced by sound waves. In biological systems, vibrations can trigger structural or biochemical changes.
Vowel: A voiced speech sound produced with an open vocal tract, resulting in periodic, lowfrequency-rich signals. Acoustically smoother and more continuous than consonants.
Zero-crossing rate (ZCR): A measure of how frequently a signal crosses zero amplitude. Higher values indicate noisy or rapidly changing signals.

